# Developmental timecourse of aptitude for motor skill learning in mouse

**DOI:** 10.1101/2024.07.19.604309

**Authors:** Taehyeon Kim, Bryan M. Hooks

## Abstract

Learning motor skills requires plasticity in the primary motor cortex (M1). But the capacity for cortical circuit plasticity varies over developmental age in sensory cortex. This study assesses the normal developmental trajectory of motor learning to assess how aptitude might vary with age. We trained mice of both sexes to run on a custom accelerating rotarod at ages from postnatal day (P) 20 to P120, tracking paw position and quantifying time to fall and changes in gait pattern. While animals of all ages were able to perform better after five training sessions, performance improved most rapidly on the first training day for mice between ages P30-60, suggesting an age with heightened plasticity. Learning this task required M1, because pharmacological inactivation of M1 prevented improvement in task performance. Paw position and gait patterns changed with learning, though differently between age groups. Successful mice learned to shift their gait from hopping to walking. Notably, this shift in gait happened earlier in the trial for forelimbs in comparison to hindlimbs. Thus, motor plasticity might more readily occur in forelimbs. Changes in gait and other kinematic parameters are an additional learning metric beyond time to fall, offering insight into how mice improve performance. Overall, these results suggest mouse motor learning has a developmental trajectory.

**Significance:** Plasticity in sensory cortex is restricted to a limited developmental window. Learning motor skills requires motor cortex plasticity, but it is unknown whether learning aptitude changes over development. Here, we define the developmental trajectory of motor learning aptitude for the accelerating rotarod task in mice, demonstrating a sensitive period for motor learning. Learning peaks at P30-P60, with mice learning to shift from hopping to walking gait to stay on the rotarod longer. Learning this task depends on M1. Further, the gait shift in forelimbs precedes hindlimbs. Knowing the peak in motor plasticity identifies the time window at which we should seek to understand the circuit basis of motor learning plasticity in cortex.

## Introduction

The critical period is a time during early development when neural circuits exhibit heightened plasticity in response to sensory experience (Hensch, 2005; Erzurumlu and Gaspar, 2012; Hooks and Chen, 2020). Studies of cortical plasticity have generally investigated sensory critical period plasticity, where manipulations at defined early ages can cause long-lasting changes in cortical connectivity and responsiveness, while plasticity is reduced or absent at later ages (Van Der Loos and Woolsey, 1973a; Knudsen and Knudsen, 1990; Gordon and Stryker, 1996). In both the visual and somatosensory systems, the timing of the critical period is defined by the ability to evoke plasticity in neuronal responses by the deprivation of the corresponding sensation, such as incoming visual or whisker stimuli (Hubel and Wiesel, 1970; Van Der Loos and Woolsey, 1973b; Katz and Shatz, 1996; Buonomano and Merzenich, 1998). For visual areas, for example, critical period plasticity is limited to ∼P19-P32 in mice (Gordon and Stryker, 1996; Hensch, 2005; Hooks and Chen, 2020). During this period, the transient removal of the incoming visual stimulus leads to drastic changes in the neuronal response, measured as a shift in ocular dominance. In contrast, the critical period in the motor system is less well-defined.

Plasticity in the primary motor cortex (M1) is vital for the learning and executing of complex motor skills (Kawai et al., 2015; Papale and Hooks, 2018; Sauerbrei et al., 2020). But few studies using animal models have characterized motor skill plasticity during developmental periods over development (Walton et al., 1992). The manipulation used to define the critical period in S1 and V1 – by eliminating incoming excitation associated with that sensory modality – is challenging to model in the motor system, as complete motor paralysis inhibits normal growth and development. In studies using motor skill acquisition, most use young adults >1 month old (Yang et al., 2009; Peters et al., 2014; Chen et al., 2015; Cichon and Gan, 2015; Adler et al., 2019). Motor skill acquisition across developmental ages in younger animals, closer to ages equivalent to the sensory critical period, is largely unexplored, particularly in ages matching the visual critical period. However, this requires addressing learning in a task that is both cortex-dependent as well as one that animals of all ages can learn. Furthermore, skill acquisition remains useful throughout life. The need to learn motor skills continues throughout life. Thus, it is worth exploring the degree to which this plasticity is altered in later ages. Investigating age-related changes in motor learning will determine whether a period of heightened plasticity for motor learning exists.

What cortical circuits might be involved? Voluntary movements and motor skill acquisition involves activity of subsets of cortical cells, task-active cells, forming local circuit ensembles that may contain from 10-50% of local neurons depending on the task (Costa et al., 2004; Dombeck et al., 2009; Hira et al., 2013; Peters et al., 2014). Motor skill acquisition is associated with structural and functional changes in these specific ensembles of excitatory neurons (Yang et al., 2009, 2014; Peters et al., 2014). Prior work has suggested changes in cortical circuit connectivity, potentially restricted to the upper cortical layers during motor learning (Jacobs and Donoghue, 1991; Kaneko et al., 1994; Rioult-Pedotti et al., 2000; Hira et al., 2013; Peters et al., 2014). This includes reweighting of long-range corticocortical and thalamocortical inputs (Kaneko et al., 1994). Thus, plasticity in upper layers is likely to be involved in motor learning. Further, inhibitory circuitry is crucial for regulating plasticity and triggering the onset and closure of critical period plasticity in sensory areas (Hensch, 2005). Similarly, reduced inhibition in motor areas is correlated with motor learning performance in normal and post-injury conditions in humans and mice (Bachtiar and Stagg, 2014; Blicher et al., 2015; Alia et al., 2016; Kida et al., 2016, 2023; Kolasinski et al., 2019). Thus, reweighting of inhibitory circuits may also play a role in regulating motor plasticity across ages. Still, whether mechanisms govern cortical plasticity for motor learning, as occurs in sensory areas, is not fully known.

To identify developmental changes in motor learning, we developed a custom accelerating rotarod task. The rotarod was made from clear plexiglass to use high-speed video to monitor changes in gait. The rod size was selected for use at any age after P15, when mice become ambulatory. Though animals of all ages learned the task, mice at ages P30-60 improved most rapidly. Learning required M1, because blocking M1 prevented improvement in the task. Tracking paw position and gait suggested differences in how the mice learned the task. This data suggests that plasticity in motor circuitry is heightened at certain ages. This should be considered when exploring circuit mechanisms for motor learning.

## Results

### Rotarod performance peaks and plateaus between P30-60

To test motor learning, we used the accelerating rotarod task, which takes advantage of walking on an accelerating rod. Mice become ambulatory by P15. Walking is part of natural locomotion that can be tracked in mice at a range of ages and sizes. Although using a skilled reaching task to explore age-dependent differences in skill learning is possible, it was not as clear that P15-P20 mice could execute this task. Thus, we preferred the rotarod, as task mice could more readily be taught. Furthermore, improvement in the task can be observed within a single training day. In the rotarod task, the time to fall and changes in foot position were measured to determine the learning curve at different ages. To monitor changes in the gait, we built a custom rotarod made of clear plexiglass. A high-speed camera was mounted at the bottom to record the behavior (Fig. 1A-C). Six age groups were used to test the age-related differences in the aptitude to learn a motor task: P15-20, P30, P45, P60, P90 and P120 (Fig. 1D-E). Behavior during the first training day for these ages tended to improve during the first half and reach a plateau within the first session, with some reduction in performance in later trials on the first training day. Improvement was measured by comparing the time to fall when the animal was at a beginner level (early phase of learning) with the time to fall when the animal reached an expert level (a performance plateau). Thus, improvement was calculated by subtracting the average time to fall measured in trials 11-13 (when the performance tends to reach plateau) from the average time to fall measured in trials 1-3 (beginner; Fig. 1D). Within the first two months of age, the improvement was greater for the animals in P30, P45, and P60 compared to those in the P15-20 group (Fig. 1E). Compared to P90-120 animals, the animals in P45-60 also showed significantly greater improvement (Fig. 1E). Notably, animals at all ages could learn the task, albeit at a slower pace (Fig. 1G).

**Figure 1.**
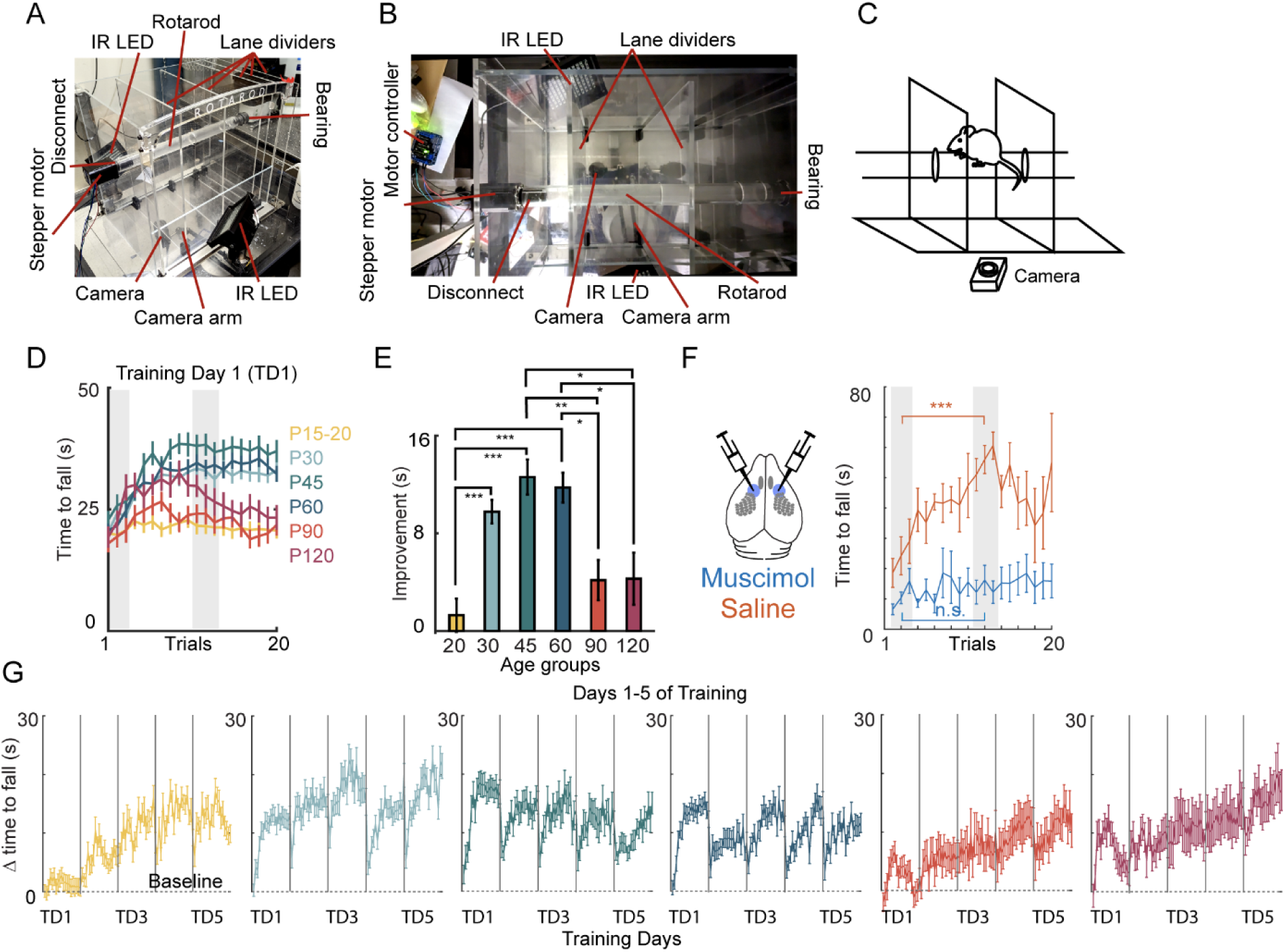
Rotarod skill acquisition plateaus around P30-P60 on training day 1. (A) Image of a clear plexiglass rotarod with high speed video. Steeper motor and camera run by computer adjacent to the operator. Multiple lanes are possible (divided by moveable lane dividers) to simultaneously run multiple mice (two lanes in the camera field of view). Infrared (IR) LEDs used for illumination during behavioral training. (B) Overhead view of the apparatus. Lane dividers opened further than during trials to illustrate capacity for sliding. (C) Schematic of rotarod task with high speed videography and barriers to create the running lane during a trial. (D) Video frame of mouse during running. Paw positions (LFP, left front paw; LHP, left hind paw; RFP, right front paw; RHP, right hind paw) and the nose and tail base are tracked using DeepLabCut. (E) Learning curve during 20 trials on training day 1 (TD1). Six colors represent the six ages tested (P15-20, P30, P45, P60, P90, and P120). Gray bars represent time windows used for early and late performance to assess TD1 improvement. (F) Improvement (in seconds) in time to fall during TD1 for six age groups. Mean+/- SEM. N=63, 92, 60 63, 27, 12. The effect of M1 inactivation by bilateral muscimol injection on rotarod skill acquisition compared to saline injected controls. N=5 (control); N=9 (muscimol). Mean+/- SEM. Significance: *, p<0.05; **, p<0.01; ***, p<0.001. (G) Improvement (in seconds) in time to fall during five training days.

To determine whether learning on the accelerating rotarod is an M1-dependent task, M1 was pharmacologically inactivated by bilaterally injecting muscimol in P30-45 animals. Mice that received bilateral injections of either muscimol or saline were permitted to recover from the procedure for 40 minutes and showed normal ambulation in the home cage after recovery. When tested on the accelerating rotarod, muscimol-injected animals did not show improvement compared to the saline-injected littermates, confirming that the task is an M1-dependent task (Fig. 1F).

Testing a behavioral task across different age groups came with some confounds. Although this task used no external rewards to exploit the natural walking behavior in mice, it is possible that age-dependent differences in the time spent walking in the home cage might contribute to rotarod performance. To assess if there was a difference in baseline locomotion, baseline motility of animals was tracked in a standard cage with infrared beam breaks to determine the number of times they crossed the length of the cage. P15 mice crossed the least number of beak breaks compared to the other age groups (Fig. 2A, p<0.001). P45 animals exhibited a greater number of beak breaks compared to P30 (Fig. 2A, p<0.001), but there were no other significant differences within these older ages.

**Figure 2.**
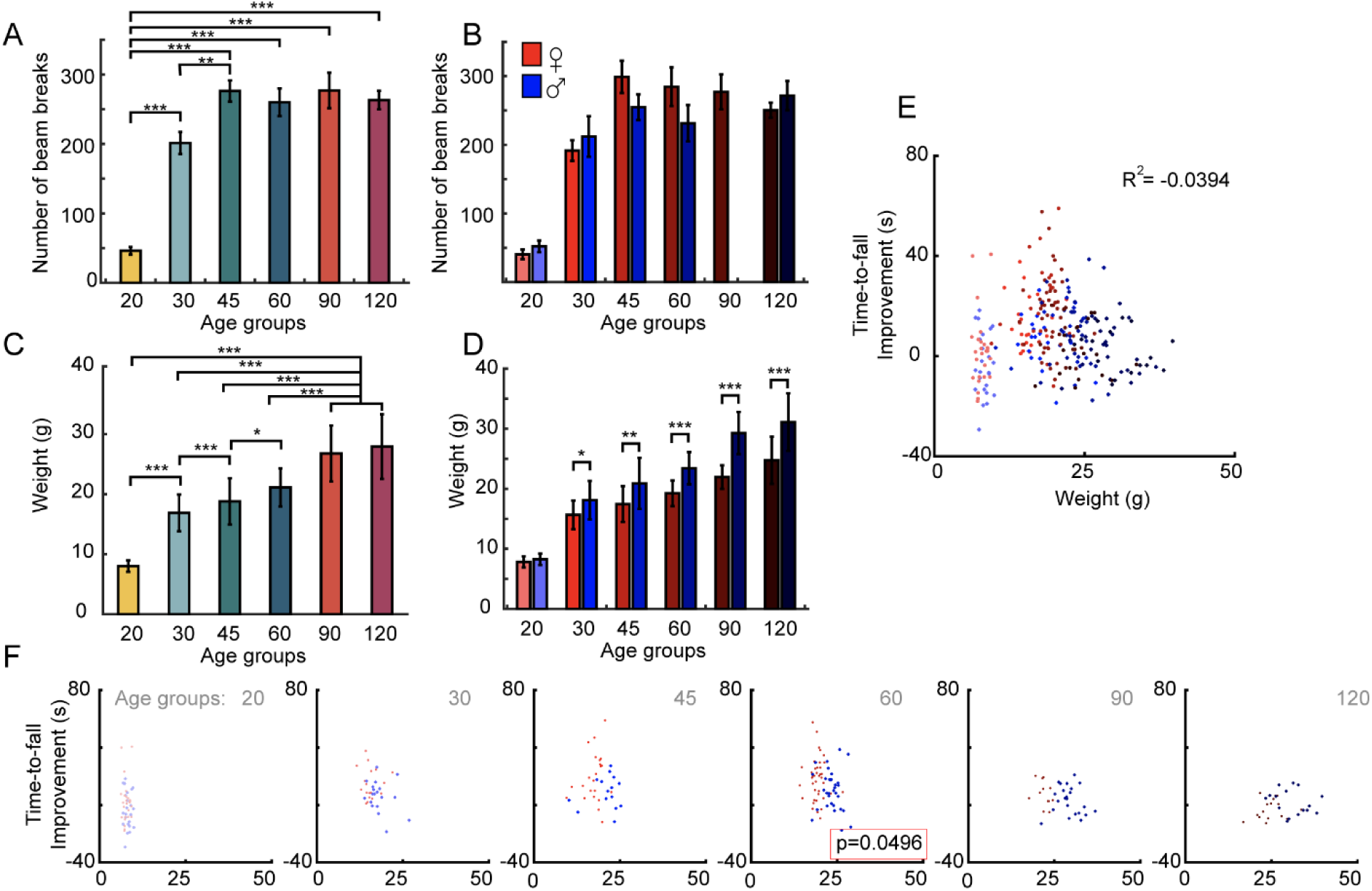
Developmental changes in mobility and weight do not account for rotarod performance. (A) Measurement of home cage walking (beam breaks) used to assess motility across developmental ages. (B) Home cage motility quantified by sex across ages. Red/blue is used to designate sex (red, female; blue, male), with brightness used to indicate age. The same color scheme is used in D-E. (C-D) Changes in weight across developmental ages, quantified for all mice or by sex. Male mice weigh more than female mice at ages older than P30. (E) Plot comparing the first training day (TD1) time-to-fall improvement versus weight. Red/blue color code is used to indicate age and sex. Linear fit is relatively poor (R^2^=-0.04). (F) Weight vs. improvement data separated by age. Significance: *, p<0.05; **, p<0.01; ***, p<0.001.

We next considered sex as a variable. Significant sex differences in improvement were observed in the P30, P45, and P60 groups (Fig. 3B, p<0.05), with young adult female mice outperforming males in these age groups. Because adult male mice tend to weigh more than female mice, we considered that the sex difference in performance might be due to weight. From P20 to P120, each age group gained significant weight (Fig. 2C). Male mice significantly outweighed female mice starting at P30 and continued to weigh more at older ages (Fig. 2D).

**Figure 3.**
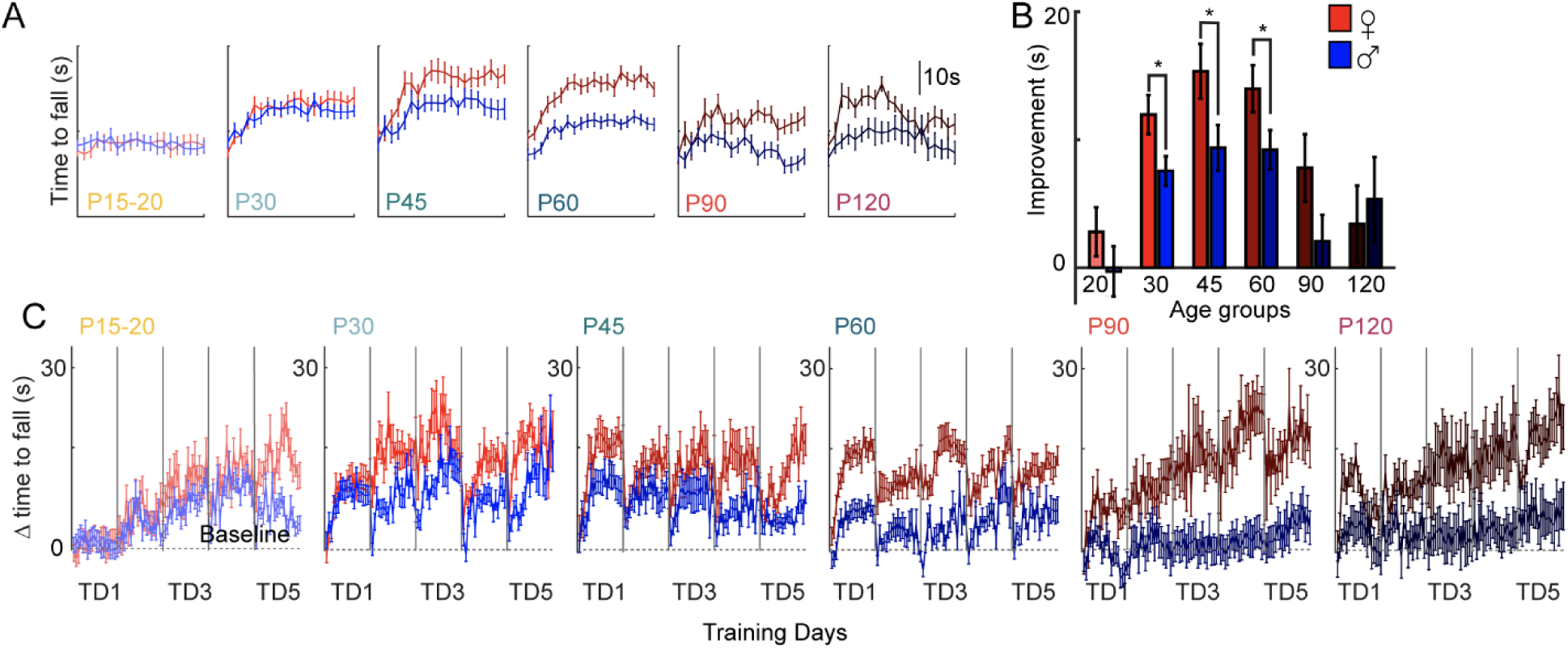
Developmental changes in rotarod performance plotted for male and female. (A) Measurement of training day 1 (TD1) performance, separated by male (blue) and female (red). (B) Summary of time-to-fall improvement, plotted as in Figure 1. (C) Measurement of rotarod performance across all training days, separated by male (blue) and female (red). Significance: *, p<0.05; **, p<0.01; ***, p<0.001.

Across all ages, there was no correlation between weight and improvement, nor was there a correlation within specific sex (Fig. 2E). P60 data alone did have a negative correlation between improvement and weight (Fig. 2F, p=0.0496). Within each sex, there was also no significant correlation of weight and performance. Therefore, weight alone did not explain differences in performance, since there were no significant differences between heavier and lighter mice at P30- P60.

### Changes in paw kinematics are correlated with better performance

Although time to fall served as an effective measure of overall task performance, a more detailed understanding of how animals improved during the task was achieved by high-speed videography to quantify changes in mouse limb movements. This approach tracked the kinematics of limb movements by acquiring a video of paw position through a clear rod. During rotarod training, four paws, the nose, and the base of the tail were tracked with DeepLabCut (Mathis et al., 2018) (Fig. 4A-D). The estimated positions of these parts were tracked in x- and y-coordinates in the video, with the x-axis aligned to the direction of rod rotation. Paw positions were filtered by likelihood (>0.95) and steps were automatically detected based on the prominence of the peak in the trajectory with a minimum interstep interval using custom software in Matlab. Step end and step start positions thus extracted were then normalized to the width of the rod (Fig. 4A-D). Step length was the normalized distance between the step end and the start (Fig. 4B). Raw position in pixels was plotted to demonstrate the accuracy of the step detection, while the paw position for all paws can be simultaneously tracked (Fig. 4C-D).

**Figure 4.**
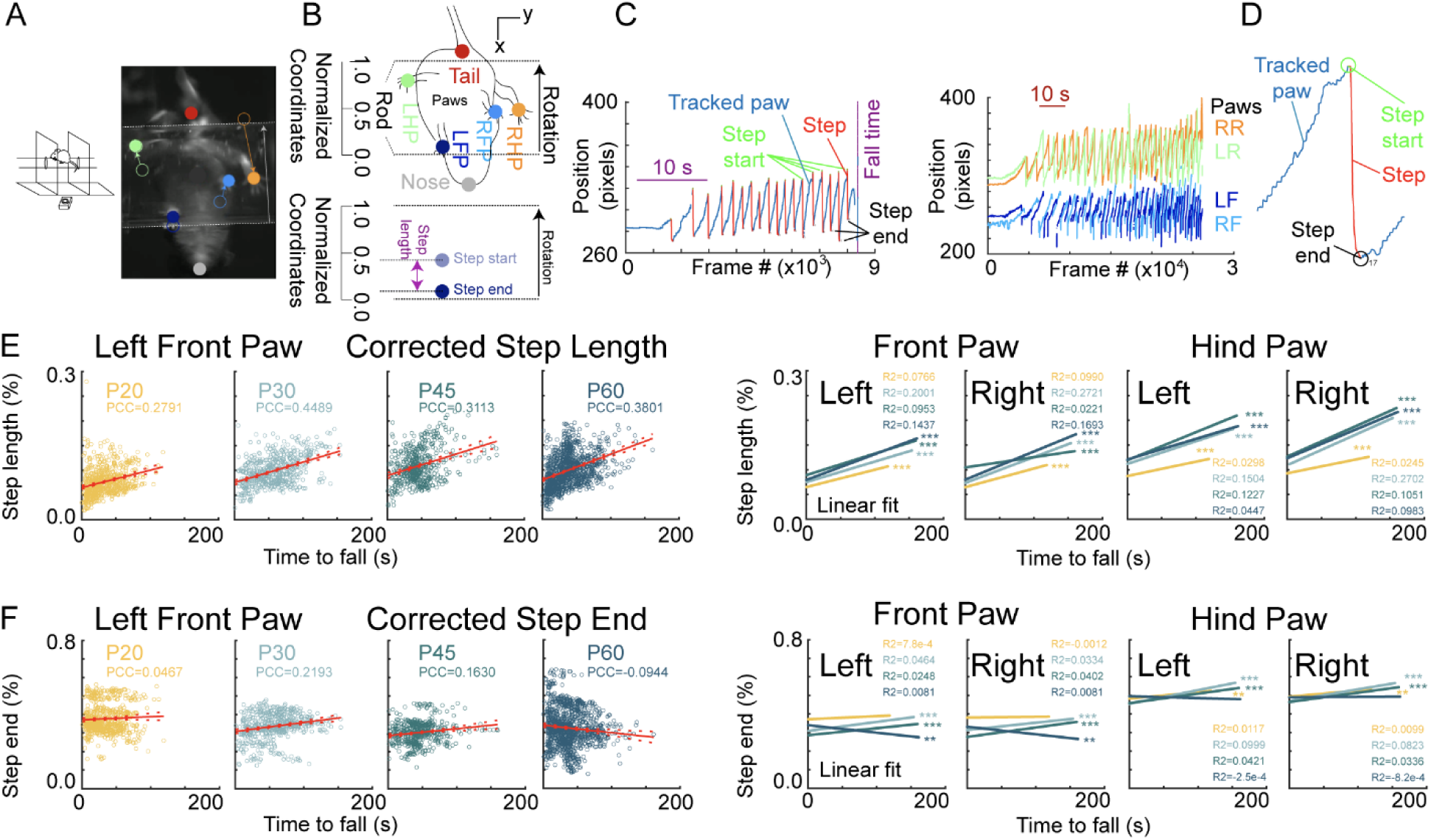
Tracking step kinematics during rotarod training. (A) Schematic of rotarod task with video frame of tracked mouse. (B) Cartoon of tracked image frame illustrating location of the mouse paws (LFP, left front paw; LHP, left hind paw; RFP, right front paw; RHP, right hind paw), the nose, and tail base using DeepLabCut. The rotarod and direction of motion are indicated, with the front and rear ends of the rod serving as 0.0 and 1.0 in normalized coordinates. At bottom, coordinates for the step start, step end, and step length are plotted to illustrate how they are determined. (C) Plot of the x-position of a single paw (LHP, left) or all paws (right) during a trial. Time given in frames (converted to seconds with example timebar). (D) Expansion of paw tracker showing step detection. Steps (red) are automatically detected (see methods). Step start (green), step end (black), and step length are then calculated. (E) Step length versus time to fall for the left front paw, plotted for individual ages (20 and above). Data are fit to a line and the linear fit compared at right. Summary plots include step length versus time to fall for the right front paw, left hind paw, and right hind paw (far right). Pearson correlation coefficients (PCC) and coefficient of determination (R^2^) are given, along with the statistical significance of the correlation (**, p<0.01, ***, p<0.001). (F) Step end point presented similarly to step length.

We then examined how paw kinematics changed during learning by comparing the normalized step length and step endpoint with performance (Fig. 4E-F), measured as time to fall. When pooling data across ages, a few general phenomena emerged. First, longer step length was associated with longer times on the rotarod. This was true across all ages measured (P20- P60). We did not image sufficient P90-P120 animals to evaluate kinematics at the oldest ages. The increase in step length was similar for both front and hind paws. Second, the step endpoint moved relative to the front edge of the rotarod. This effect was strongest for front paws but was present to a lesser degree for hind paws (as expected if step size also increases). This effect, however, varied with developmental age. In P60 animals, the step end point shifted closer to the front end of the rod. At other ages, the step end position is static (P20) or moves away from the edge (P30 and P45). This may be due to differences in animal size relative to the rod. This effect is similar for both left and right forepaws (Fig. 4F right).

### Changes in gait differ between forelimb and hindlimb

The rotarod task involves not only kinematic changes in individual paws, but also the coordination of left and right paws. Animals engaged in different types of gait patterns during the task – simultaneously moving both paws (hopping; Fig. 5A) or alternating the left and right paws (walking; Fig. 5A). To determine how the gait pattern changed, we needed to consider the phases of steps between left and right paws. We compared the step times of the paws and assigned each step a score (range 0.0 to 1.0). The step time of the left paw (Fig. 5C, green for the left hind paw) was marked. The time between the two left steps was normalized from 0.0 to 1.0. The time of the right hind paw step (orange) was then computed as a fraction of the relative distance between the left steps. Alternating gait occurred when the phase difference of the left and right paws was closer to 0.5 while the simultaneous or hopping gait was closer to 0.0 or 1.0 (Fig. 5C). Note the color map at the bottom which signifies the alternating gait as red and the hopping gait as blue. The scale was reflected because we treated step phase scores from 0.5 to 1.0 as equivalent to 0.5 to 0.0 to assess alternating gait. To determine changes in gait pattern on the first training day and how these differed with limb, age, and trial number, the data were plotted as follows. For each step taken, such as the first step on the first trial (20), the mean step phase score was computed across all mice tracked, and the corresponding color was plotted as a tick mark (Fig. 5D). These were generally blue (simultaneous/hopping gait). This was repeated for each subsequent step and for each trial. The number of steps included in the average varied across the row for each trial, reflecting the variability in performance (as observed in time to fall) within and across trials. Qualitatively, the gait pattern generally shifts from hopping to walking (alternating gait) for both front limb and hind limb. This occurs to some degree for all ages tested. This shift seemed to occur more rapidly for the front paws than the hind paws, suggesting that the transition to a walking gait can occur at different times for the two pairs of limbs.

**Figure 5.**
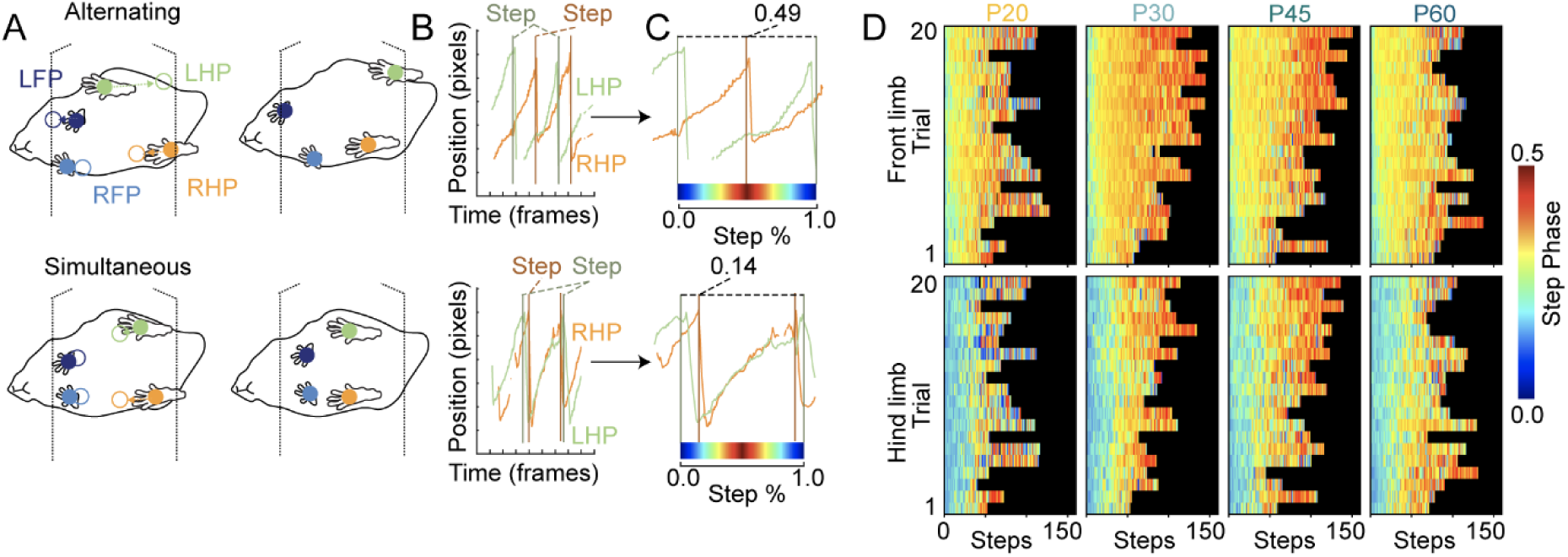
Gait pattern changes with age and training. (A) Schematic of paw tracking during alternating gait/walking (top) or simultaneous gait/hopping (bottom), with front (LFP, RFP) and hind (LHP, RHP) paws tracked. (B) Using the hind paws to illustrate, LHP (green) and RHP (steps are shown, with vertical lines incidating step start times. (C) The step start times for one foot are scaled from 0.0 (when the first LHP step begins) to 1.0 (when the next LHP step begins) and the timing of the RHP step assigned a value from 0.0 to 1.0 (in the top example, 0.49). The color map at bottom indicates the color to be used in the subsequent plot to illustrate phase (with red representing 0.5/alternation and blue for 0.0/simultaneity). The step phase map is mirrored for values ranging from 0.5-1.0. (D) Overall gait pattern is plotted by limb (front limb, top; hind limb, bottom) and age (from left to right, panels for P20 to P60). Within each panel, the rows represent trials on training day 1 with the bottom row as trial 1 and the top row as trial 20. For each step, starting with the first step taken, step phase is plotted as the average across all mice in that condition. The number of steps included in the average thus varied across the row for each trial (when some mice fall off the rod).

To determine whether the gait pattern shifted significantly on the first training day, the cumulative frequencies of the step phases at four different stages of the training were plotted (Fig. 6A): beginner (trials 1-5), early learning (trials 6-10), mid-learning (trials 11-15), and late learning (trials 16-20). The offset in the step phase between front limbs (dotted lines, lower right) and hind limbs (solid lines, upper left) was evident in the data, suggesting that the front and hindlimbs employ different strategies during rotarod training. Cumulative frequencies of the phases in the beginner stage were compared to the other three stages. Across all age groups, the front paws shifted to alternating gait starting from early learning at all ages (trials 1-5 compared to trials 6- 10, 11-15, and 16-20) (Fig. 6A, left). On the contrary, the shift from simultaneous hopping to alternating in the hind paws was delayed in an age-dependent manner (Fig. 5D and 6B). P20 groups started alternating hind paws by late learning (trials 16-20); P30 animals by mid-leaning; P45 and P60 groups by early learning (trials 6-10).

**Figure 6.**
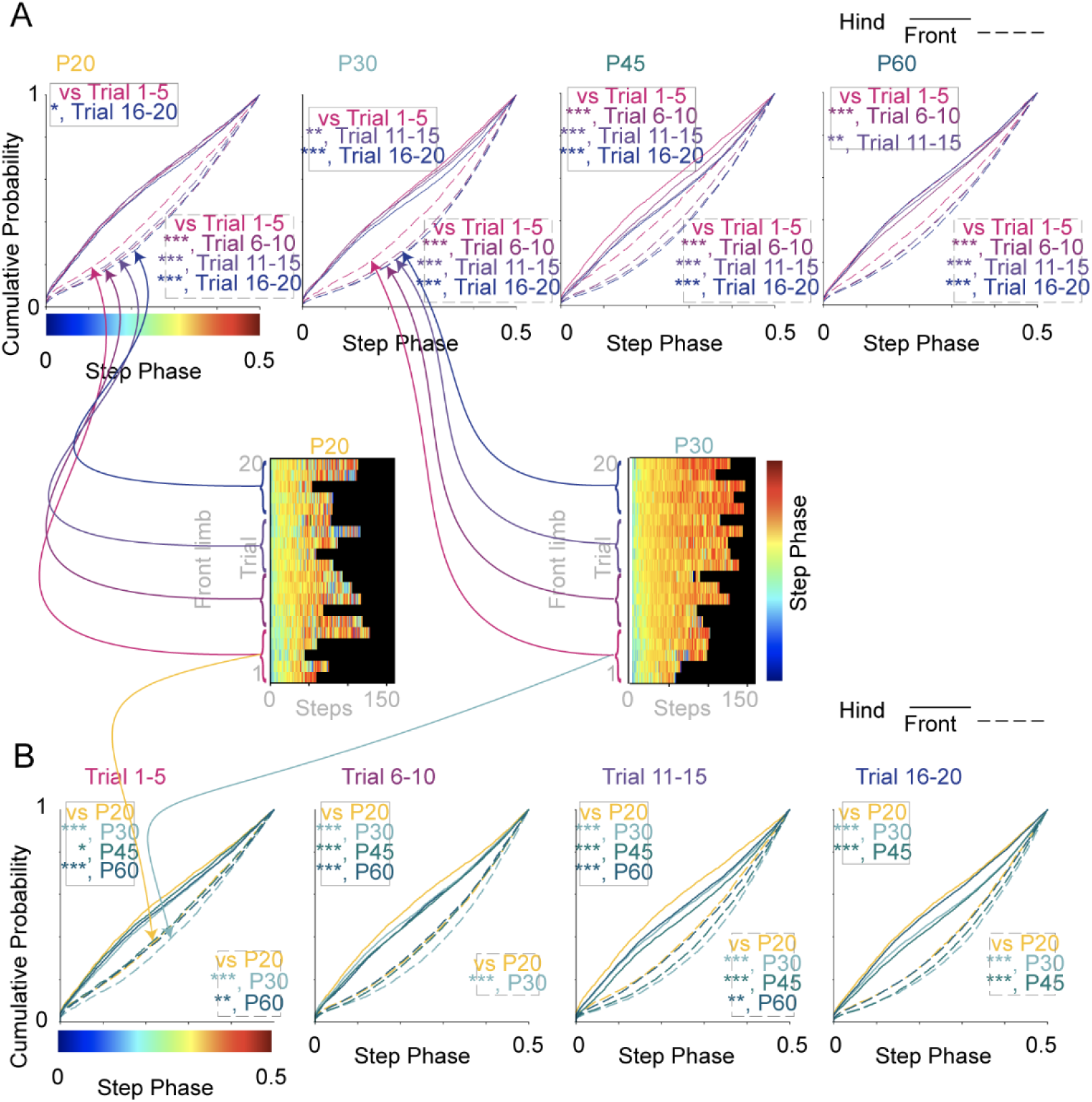
Statistical comparison of gait pattern changes across trials and ages. (A) Cumulative probability plots for the data in Fig. 5D to compare step phase across trial number (in groups of Trials 1-5, 6-10, 11-15, and 16-20). Dashed line represents front paws; solid line represents hind paws. (B) Cumulative probability plots for the data in Fig. 5D to compare step phase across age (in groups of P20, P30, P45, and P60). Dashed line represents front paws; solid line represents hind paws. Cartoon illustrates the categories from which the individual step data is drawn to make the cumulative probability plot. The step data is averaged across mice in Fig. 5D, but here each individual step phase value is used. Significance: *, p<0.05; **, p<0.01; ***, p<0.001.

However, the change in the distribution of the gait patterns over time within a trial was not well-quantified in this manner. We wanted to compare whether there was a difference in the emergence of a gait pattern and compare between age groups and trials. But this was complicated by the fact that trials differed in number of steps. To compare this fairly across trials, the entire series of steps in a trial was pooled and averaged into deciles (nine bins) which were then compared between deciles, trials, and age groups (Fig. 7A). This appeared similar to the raw data (Fig. 7D), with all trials now scaled to the same length. Qualitatively speaking, the first decile of the front paws tends to be more hopping (in blue) and changes into more alternating (in orange). On the contrary, the deciles of the hindpaws seem to be less alternating than the front limbs (in blue to yellow). This approach allowed us to look at the changes in gait patterns despite the difference in the number of steps in each trial (Fig. 7A).

**Figure 7.**
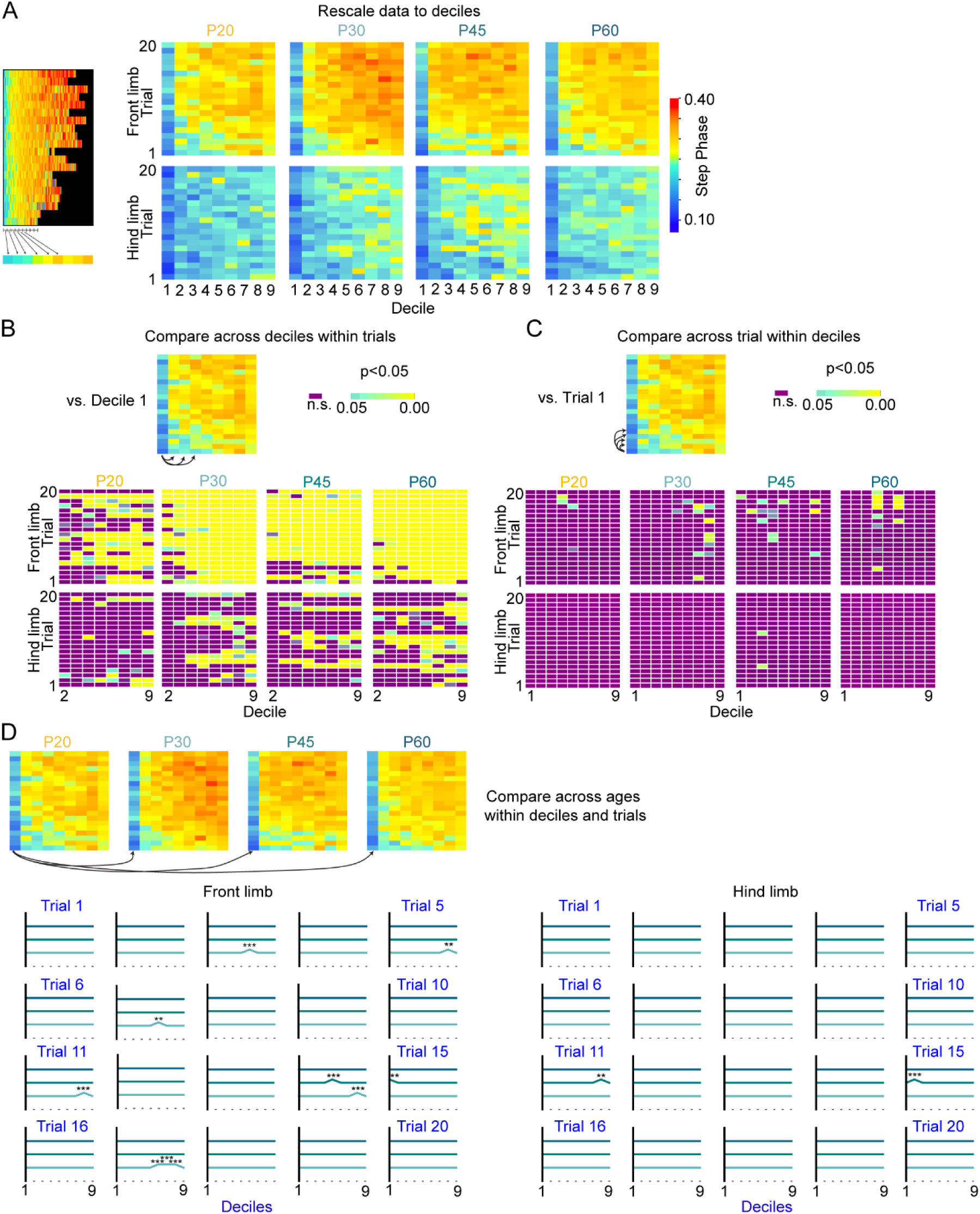
Visual representations of p-values for the gait analysis. (A) Due to variations in the number of steps, the number of steps in a trial were divided into deciles to facilitate the comparison of average step phases across trials and age groups. The average step phase value for each age for front and hind limbs is plotted. (B) The P-value plot of deciles compares the first decile of trials 1-20 to rest of the deciles (2-9) for each trial. The top row compares front paws at four ages and the bottom row compares hind paws. Non-significant p- values are purple; p-values < 0.05 are as shown in the lookup table. (C) The P-value plot comparing across trials compares each decile from trials 2-20 with the corresponding decile from trial 1. The top row compares front paws at four ages and the bottom row compares hind paws. For instance, the front paw in P20 Decile 3 from trial 3 is compared to Decile 3 from trial 1. Non- significant p-values are purple; p-values < 0.05 are as shown in the lookup table. (D) The P-value plot of deciles compared across ages assess P20 versus P30, P45, and P60 groups for 20 trials. Each individual graph represents the comparison between the four age groups for a given trial. The X-axis denotes deciles, with a specific trial’s decile being compared to the corresponding decile of the P20 group. For example, the front paw in P30 Decile 9 is compared to P20 Decile 9 in the trial 1 graph, and if significantly different, it is indicated by a bump in the y-axis. Significance: *; p<0.05. **;p<0.01, ***;p<0.001.

First, to compare whether and how the gait pattern changed during a trial, we compared the step phase at the beginning of the trial to the step phase at later time points. The step phase at the beginning was quantified as the step phase values for the first decile of a given trial (the first ∼10% of steps) to the rest of the deciles in the same trial (the subsequent steps in ∼10% blocks; Fig. 7B, arrows). This showed the strongest effect: the phase of the front paws, particularly in P30-60 groups, showed significant differences compared to the first deciles across many trials. The difference in the phase of the front paws (Fig. 7B, top) during the first decile compared to later deciles is present as early as the first trial. This effect progressively becomes more prominent with training as even early deciles prove different than the first decile in later trials. Hindpaws seemed to show few significant differences across deciles (Fig. 7B, bottom). Animals in P30-P60 showed some difference in mid to late deciles mostly in early to mid-learning phase, while the P20 group showed minimal difference in the phase of hind paws.

Second, to test whether there was a difference in gait across trials, the decile of the first trial was compared to the corresponding decile of the rest of the trials (trials 2-20, see arrows). This effect was considerably weaker with fewer significant comparisons). In the P20 group, only front paws showed a small difference in the late learning phase (Fig. 7C, top). In the P30 group, the difference in front paw occurred by early-to-late learning (as early as trial 5) in late deciles (Fig. 7C, top). In the P45 group, the difference in the front paw appeared later and became more evident around mid-learning, particularly in early-mid deciles. In the P60 group, differences in front paw started to predominantly appear in late trials during late learning (Fig. 7C). As before, the difference in hind paw gait across the age groups was minimal, only observed in P45 (Fig. 7C).

Lastly, to compare explore age-related differences across age groups, we compared a decile in a given trial of P20 to a corresponding decile in the same trial of P30, P45 or P60 (Fig. 7D, long arrows). Front paws of P30 started to show differences in early trials and differed from P20 in some trials (Fig. 7D). On the other hand, P45 and P60 showed a difference to P20 during mid-learning (Fig. 7D). Hind paws gait patterns only differed in mid-learning in P45 group (Fig. 7D).

## Discussion

### Rotarod motor skill learning is fastest in a developmental window

To address age-dependent differences in motor learning, we first used a walking task that was consistent with the range of natural behaviors in mice. Walking is a common behavior in vertebrates that relies, in part, on central pattern generators in the spinal cord (Grillner and Wallén, 1985). For mice, walking begins naturally prior to the ages we test. To assess whether there is a sensitive (critical) period in the ability to acquire a motor skill, we tested mice as young as P15 (an age at which they are able to execute the task) up to age P120 on the accelerating rotarod. Using time to fall as a measure of learning, these data show that first-day improvement in performance on the rotarod varies with age, plateauing at P30-60. This suggests that there are differences in the aptitude for motor learning across developmental time. This is unlikely to be due to a difference in the baseline locomotion, which does not show selectively heightened activity at P30-60 (Fig. 2A-B). Previous experiments had also shown that prior exposure to exercise did not affect learning (Buitrago et al., 2004). Control experiments also ruled out the effects of weight (Fig. 2E). Since improvement does occur over five days of training for all age groups of mice (Fig. 2F), the animals are physically capable of executing the task at all ages.

The rotarod task is not associated with explicit reward, as mice walk naturally and running is an intrinsically motivated behavior (Sherwin, 1998; Greenwood and Fleshner, 2019). All ages can learn: some individual mice learn faster (within the first training day), while others require more training. However, it is not clear how to distinguish whether motivational state differs across ages and might contribute to task performance. If the motivational state is the confounding factor driving the age-related difference in the performance on the rotarod, it suggests that age groups that performed poorly on the first training day (P20, P90 and P120) had low motivational state but were able to regain/increase the motivation on the following training days 2-5 to show improvement measured in time to fall, with a sufficient motivational swing as to influence the training performance on a day-to-day basis. This magnitude of fluctuation in motivation seems unlikely to explain the results. It is possible to speculate that other factors, such as anxiety (from being placed on an elevated rod), contribute to performance differences at different ages. An assay for fear might clarify this, though a nonlinear age-dependent variation in fear would be interesting to explore in other circuits.

A second aspect that is also hard to disambiguate is whether learning in mice occurs with understanding the goal of the task (walk so as to stay on the rod) becomes clear, or whether the goal is clear and learning is a shift in precise control of movement. Both shifts could result in a successful increase in time to fall. Although these experiments do not clearly distinguish between these alternatives, tracking the kinematics of paw movement suggests that, at least in part, better performance is associated with changes in motor control. This seems to be the case because the step lengths for all paws evolve during the task (Fig. 4E), front paw step endpoints are targeted closer to the end of the rod, at least for P60 (Fig. 4F), and the gait pattern evolves with time during the task (Fig. 6 and 7B).

Because spinal circuits also play a role in the control of walking, it is important to test whether rotarod skill learning depends on the motor cortex. This task has been used in previous studies of the motor cortex to assess circuit changes during learning. Specifically, rates of spine formation and retention in forelimb M1 change during rotarod learning (Yang et al., 2009, 2014). Not only are there structural changes, but the firing patterns of neural ensembles show task-specific modulation during training and reorganize over several sessions to support motor learning (Costa et al., 2004). Together these imply that circuit changes assist in acquiring a new skill. Consistent with this, our data shows that bilateral inhibition of forelimb M1 with muscimol during rotarod training blocks motor learning relative to saline-injected control mice (Fig. 1F). Normal walking (in the home cage), however, was not affected in muscimol-treated mice and these mice exhibited other non-learned motor behaviors, such as jumping. Collectively, this suggests a necessary role for forelimb M1 in rotarod learning.

### The transition from hopping to walking gait facilitates motor learning best in P30-P60 mice

Markerless tracking (Mathis et al., 2018) enabled tracking paw position and segmenting paw trajectories into steps (Fig. 4). This was essential to understanding changes in paw kinematics as well as comparing steps to understand gait changes. Step length was tracked for all four paws and showed a consistent effect across age groups. For both forepaw and hindpaw, steps became longer as mice were more successful at remaining on the rotarod. Further, while the trend was similar, P20 mice took shorter steps compared to the other age groups, consistent with smaller body size. Targeting of the step endpoint relative to the end of the rod was more complex. In general, the endpoint location was relatively stable during learning across ages, with the exception of the P60 mice. P60 animals developed a strategy of placing the forelimb step closer to the front edge of the rotarod. The effect was smaller but present for the hind limb because the step length became longer. This suggested that mice were able to change some of the kinematic parameters of their steps, including targeting a specific location on the rod. Intracortical somatosensory information plays an important role in motor learning and may be implicated in emergence of the kinematic patterns (Chang et al., 2022; Huang et al., 2024). Whether this step location targeting requires visual feedback is not clear. The experiments were run in the dark with infrared light, and thus the mice likely had no visual feedback during the task.

More interesting is when individual steps are analyzed for changes in the gait pattern. Overall, the gait shifts from hopping to walking as individual trials proceed. But the shift is more pronounced for certain trials and at certain ages. Further, the shift from hopping to walking appears to happen at different times for forelimbs and hindlimbs, suggesting that the circuitry controlling the mode of locomotion can operate independently for each pair of limbs. The shift from a hopping to a walking strategy enables animals to stay on the rod, though it is unclear whether a split in gait pattern between forelimb and hindlimb would occur in other more naturalistic conditions. Notably, humans are able to walk at different speeds between the left and right leg on a split treadmill (Iturralde and Torres-Oviedo, 2019), but interpreting this adaptation is difficult. That the targeting of the paws closer to the front edge of the rod and the change in gait occurs in a more pronounced way for the forelimb suggests that motor learning may involve this limb more than the hindlimb. If the learning were simply a response to the movement of the rod, sensory feedback, and balance, this might be more likely to involve both pairs of limbs, while specific involvement of the forelimbs might imply a more deliberative or conscious effort that involves the frontal cortex.

What about differences in the gait shift? These data can be compared across bins within the same trial, for the same bin across trials, and across ages. Within the first training day, the forelimb step phase shifts significantly towards walking for ages after trial 5 (Fig. 6A). However, the early shift for hindlimbs in the same comparisons is strongest only for P45 (Fig. 6A). This suggests that forelimb performance is more plastic in this assay, and that the hindlimb plasticity is also enhanced at certain ages (P45). When comparing across ages at the same trial number, older animal groups perform better than P20 animals (Fig. 6B). This shift appears larger and has more statistical support for mid-learning trials, where the improvement tends to plateau (Fig. 6B, trials 11-15). These plots, however, do not account perfectly for shifts in gait since they compare steps across the entire trial (but see Fig. 7B). Subdividing the steps into bins by a fraction of steps taken on a given trial, a more detailed comparison with fewer steps per bin can be made, though this reduced the statistical power. In this approach, the strongest effects were observed across deciles in a given trial for the front limbs, further supporting the plasticity of the front limb.

Another potentially interesting reason for different engagement of the forelimb versus hindlimb is that there may be different needs for the forelimb vs. hindlimb. Many other motor learning tasks oftentimes involve forelimb-only use tasks. Rodents are capable of making very dexterous manipulations using their forepaws. Thus, it is consistent with the idea that forelimbs may exhibit more plasticity during motor learning. In addition, anatomical data seem to show the difference in reciprocal connectivity between S1 and M1 depending on the location in the somatotopic map (Zingg et al., 2014). More projections between forelimb S1 and M1 are observed compared to the projections between hindlimb S1 and M1 (Zingg et al., 2014). Silencing of S1 not only interfered with locomotion but also disrupted a motor learning skill, suggesting a role for somatosensory input in learning (Karadimas et al., 2019; Chang et al., 2022; Huang et al., 2024).

Our study shows that with a motor learning paradigm that requires all four limbs, there is a sensitive period during which animals show better aptitude for learning the rotarod task. This suggests a specific age window in which circuitry should be explored to better understand the circuit mechanisms underlying motor system plasticity. Notably, while P30-P60 mice are not fully mature, this window is later than the critical period for visual cortex in mice, which occurs about P19-P32 (Gordon and Stryker, 1996). The motor learning window also does not fully close as traditional critical period plasticity for sensory areas does. Of course, the nature of these circuit changes and the cell types involved may continue to vary with age. Further, because plasticity associated with learning may correspond to a change which may leave an anatomical trace, it is possible traces of former plasticity may be retained (Yang et al., 2009). Thus, it will be of interest to further ask whether a brief exposure to the task during the youngest age (P20) will have a lasting impact on performance at later ages (P60-90), which did not show drastic improvement on the first day. Though shifted to a slightly later time in development, it will be interesting understand how the long-term effects of early motor plasticity compare to the anatomical and functional traces left by early plasticity in sensory systems.

## Materials and Methods

### Mice and Behavior

Animals were trained on an accelerating rotarod. The rotation accelerated uniformly from 0 to 90 RPM in 3 minutes. Animals performed 20 trials in a session and one session per day. A subset of animals (those that were not sacrificed for brain slice experiments after the first training day) continued to train for 5 consecutive days. The custom-designed rotarod (Fig. 1) was built from clear plexiglass. A stepper motor (uxcell) was connected to a clear rod (20 mm diameter) and was controlled by an Arduino board. Panels dividing the lanes of the rotarod were able to slide along the rod to adjust the width of the running lane to the size of the animal running on the rotarod. At the bottom, a high-speed camera (FLIR Blackfly S USB3) was mounted to record the animal during the task. Six age groups were tested: postnatal day(P) 15-20 (N=92), P30 (N=114), P45 (N=87), P60 (N=125), P90 (N=33), and P120 (N=27). Time to fall was measured throughout all trials. The animal’s performance was recorded using the high-speed camera at 200 frames per second. The youngest age group includes P15-20. By P15, animals have exhibited mature levels of posture and reflexes (Dupuis et al., 2024). However, even though animals become ambulatory by P15 and can stay on the rotarod, this age was not ideal for rotarod training for future experiments involving skull mounted devices, because the skull is not mature and the animals are not weaned from their parents. Therefore, P20 animals were more suitable to test in both the rotarod task for behavior and brain slice electrophysiology. To measure the baseline locomotion, a custom-built infrared beam-breaking system was used. In a clear standard mouse holding cage, two pairs of infrared (IR) transmitters and receivers (Adafruit) were mounted onto the side. Mice were tested in the same age groups as the rotarod test; P15-20 (N=30), P30 (N=34), P45 (N=35), P60 (N=26), P90 (N=5), and P120 (N=15). Animals were placed one at a time in the cage and allowed to roam freely within the cage for 5 minutes. Using Arduino, the number of beam breaks of each beam was recorded when the animal broke IR beams.

### Behavioral Analysis

Improvement measured in time to fall was calculated by taking the average of trials 1-3 and subtracting it from the average of trials 11-13 (Fig. 1D). A one-way ANOVA was performed to compare the improvement in six age groups. Tukey’s multiple comparison test was used after the ANOVA. For single comparisons (muscimol), t-test was used. The same analysis was used to compare the baseline locomotion. To determine the difference in weights between males and females within an age group, t-test was used. Correlation coefficient was computed to determine whether there is a significant correlation between weights and the improvement with corrected alpha value using Bonferroni correction.

We focused on P20-60 animals, which correspond to the ages of the animals recorded for slice electrophysiology. The body positions of the animals were estimated using DeepLabCut (Mathis et al., 2018). The nose, right front paw, left front paw, right hind paw, left hind paw and base of the tail were annotated manually during the model training and subsequently tracked for analysis. Estimated position data points with likelihood of less than 0.95 were excluded from the analysis. To analyze the changes in the position of a single paw, steps were detected using custom software. The start and end of a step were converted to location relative to the front (0.0) and rear (1.0) edge of the rotarod (Fig. 4). This normalized measure is thus referred to as corrected step length, (in contrast to the raw step length in pixels). The step length was calculated by measuring the distance between the step start and end. To analyze the changes in the gait pattern, the phase of the left and right paws (Fig. 5) was determined by comparing the location and timing of one step between the corresponding steps with the opposite paw, for example where a left hind paw (LHP) step occurred between two right hind paw (RHP) steps (Fig. 5A-C). When the gait patterns were averaged across deciles (Fig. 7), trials in which animals had taken less than 10 steps were excluded, as the number of steps was insufficient. When the number of steps cannot be evenly distributed across the deciles, the remainder steps were averaged in the last decile. For the gait analysis, one-way ANOVA was performed to compare the gait phases between the deciles and the trials. Tukey’s multiple comparison test was used after the ANOVA. Comparison between a decile and trial of P20 to the corresponding decile of P30, P45, and P60 were made using t-test with corrected alpha value using Bonferroni correction.

### Muscimol Injection

To determine whether M1 is necessary to learn the accelerating rotarod task, M1 was pharmacologically inactivated. Muscimol (5 mg/ml, 40 nL per hemisphere) or saline was injected into forelimb M1 bilaterally in animals P30-45 (600 µm anterior, 1500 µm lateral, 500 and 800 µm depth, with 20 nL at each depth). 40 minutes following removal from anesthesia, the animals were tested on the accelerating rotarod. Experiments were performed blind to the injection condition (muscimol vs. saline).

## Acknowledgements

We thank Quincy Erickson-Oberg for assistance with the beam break experiment, Tom Deakin for assistance with rotarod experiment, and Kyle Bannerman for assistance with the gait analysis. We thank Omar Gharbawie, Chinfei Chen, Nuo Li, Alison Barth, Simon Chen, Caroline Runyan, Ghanshyam Sinha, Mariah Berchulski, and other members of the Hooks lab for feedback on the manuscript. This work was supported by a NARSAD Young Investigator Award (BMH), NIH NINDS R01 NS103993 (BMH), a Whitehall Foundation award (BMH), and an American Heart Association Predoctoral Fellowship (TK).

